# Structural basis for broad coronavirus neutralization

**DOI:** 10.1101/2020.12.29.424482

**Authors:** Maximilian M. Sauer, M. Alexandra Tortorici, Young-Jun Park, Alexandra C. Walls, Leah Homad, Oliver Acton, John Bowen, Chunyan Wang, Xiaoli Xiong, Willem de van der Schueren, Joel Quispe, Benjamin G. Hoffstrom, Berend-Jan Bosch, Andrew T. McGuire, David Veesler

## Abstract

Three highly pathogenic β-coronaviruses crossed the animal-to-human species barrier in the past two decades: SARS-CoV, MERS-CoV and SARS-CoV-2. SARS-CoV-2 has infected more than 64 million people worldwide, claimed over 1.4 million lives and is responsible for the ongoing COVID-19 pandemic. We isolated a monoclonal antibody, termed B6, cross-reacting with eight β-coronavirus spike glycoproteins, including all five human-infecting β-coronaviruses, and broadly inhibiting entry of pseudotyped viruses from two coronavirus lineages. Cryo-electron microscopy and X-ray crystallography characterization reveal that B6 binds to a conserved cryptic epitope located in the fusion machinery and indicate that antibody binding sterically interferes with spike conformational changes leading to membrane fusion. Our data provide a structural framework explaining B6 cross-reactivity with β-coronaviruses from three lineages along with proof-of-concept for antibody-mediated broad coronavirus neutralization elicited through vaccination. This study unveils an unexpected target for next-generation structure-guided design of a pan-coronavirus vaccine.

## Introduction

Four coronaviruses mainly associated with common cold-like symptoms are endemic in humans, namely OC43, HKU1, NL63 and 229E, whereas three highly pathogenic zoonotic coronaviruses emerged in the past two decades leading to epidemics and a pandemic. Severe acute respiratory syndrome coronavirus (SARS-CoV) was discovered in the Guangdong province of China in 2002 and spread to five continents through air travel routes, infecting 8,098 people and causing 774 deaths, with no cases reported after 2004(Drosten et al., 2003; Ksiazek et al., 2003). In 2012, Middle-East respiratory syndrome coronavirus (MERS-CoV) emerged in the Arabian Peninsula, where it still circulates, and was exported to 27 countries, infecting a total of ∼2,494 individuals and claiming 858 lives as of January 2020 according to the World Health Organization(Zaki et al., 2012). A recent study further suggested that undetected zoonotic MERS-CoV transmissions are currently occurring in Africa(Mok et al., 2020). A novel coronavirus, named SARS-CoV-2, was associated with an outbreak of severe pneumonia in the Hubei Province of China at the end of 2019 and has since infected over 64 million people and claimed more than 1.4 million lives worldwide during the ongoing COVID-19 pandemic(Zhou et al., 2020; Zhu et al., 2020b).

SARS-CoV and SARS-CoV-2 likely originated in bats(Ge et al., 2013; Hu et al., 2017; Li et al., 2005; Yang et al., 2015; Zhou et al., 2020) with masked palm civets and racoon dogs acting as intermediate amplifying and transmitting hosts for SARS-CoV(Guan et al., 2003; Kan et al., 2005; Wang et al., 2005). Although MERS-CoV was also suggested to have originated in bats, repeated zoonotic transmissions occurred from dromedary camels(Haagmans et al., 2014; Memish et al., 2013). The identification of numerous coronaviruses in bats, including viruses related to SARS-CoV-2, SARS-CoV and MERS-CoV, along with evidence of spillovers of SARS-CoV-like viruses to humans strongly indicate that future coronavirus emergence events will continue to occur(Anthony et al., 2017; Ge et al., 2013; Hu et al., 2017; Li et al., 2019; Li et al., 2005; Menachery et al., 2015; Menachery et al., 2016; Wang et al., 2018; Yang et al., 2015; Zhou et al., 2020).

The coronavirus spike (S) glycoprotein mediates entry into host cells and comprises two functional subunits mediating attachment to host receptors (S_1_ subunit) and membrane fusion (S_2_ subunit)(Ke et al., 2020; Kirchdoerfer et al., 2016; Turoňová et al., 2020; Walls et al., 2020b; Walls et al., 2016a; Walls et al., 2017; Wrapp et al., 2020). As the S homotrimer is prominently exposed at the viral surface and is the main target of neutralizing antibodies (Abs), it is a focus of therapeutic and vaccine design efforts(Tortorici and Veesler, 2019). We previously showed that the SARS-CoV-2 receptor-binding domain (RBD, part of the S_1_ subunit) is immunodominant, comprises multiple distinct antigenic sites, and is the target of 90% of the neutralizing activity present in COVID-19 convalescent plasma(Piccoli et al., 2020). Accordingly, monoclonal Abs (mAbs) with potent neutralizing activity were identified against the SARS-CoV-2, SARS-CoV and MERS-CoV RBDs and shown to protect against viral challenge *in* vivo (Alsoussi et al., 2020; Barnes et al., 2020a; Barnes et al., 2020b; Brouwer et al., 2020; Corti et al., 2015; Hansen et al., 2020; Hassan et al., 2020a; Liu et al., 2020; Piccoli et al., 2020; Pinto et al., 2020; Rockx et al., 2008; Rockx et al., 2010; Rogers et al., 2020; Seydoux et al., 2020; Tortorici et al., 2020; Walls et al., 2019; Wang et al., 2020a; Zost et al., 2020). The isolation of S309 from a recovered SARS-CoV individual which neutralizes SARS-CoV-2 and SARS-CoV through recognition of a conserved RBD epitope demonstrated that potent neutralizing mAbs could inhibit β-coronaviruses belonging to different lineage B (sarbecovirus) clades (Pinto et al., 2020). An optimized version of S309 is currently under evaluation in phase 3 clinical trials in the US. Whereas a few other SARS-CoV-2 cross-reactive mAbs have been identified from either SARS-CoV convalescent sera (Huo et al., 2020; ter Meulen et al., 2006; Wec et al., 2020; Yuan et al., 2020) or immunization of transgenic mice (Wang et al., 2020a), the vast majority of SARS-CoV-2 S-specific mAbs isolated exhibit narrow binding specificity and neutralization breadth.

Although the COVID-19 pandemic has accelerated the development of SARS-CoV-2 vaccines at an unprecedented pace(Case et al., 2020; Corbett et al., 2020; Folegatti et al., 2020; Hassan et al., 2020b; Jackson et al., 2020; Mulligan et al., 2020; Sahin et al., 2020; Walls et al., 2020a; Yu et al., 2020; Zhu et al., 2020a), worldwide deployment to achieve community protection is expected to take many more months. Based on available data, it appears unlikely that infection or vaccination will provide durable pan-coronavirus protection due to the immunodominance of the RBD and waning of Ab responses, leaving the human population vulnerable to the emergence of genetically distinct coronaviruses(Edridge et al., 2020; Piccoli et al., 2020). The availability of mAbs and other reagents cross-reacting with and broadly neutralizing distantly related coronaviruses is key for pandemic preparedness to enable detection, prophylaxis and therapy against zoonotic pathogens that might emerge in the future.

We report the isolation of a mAb cross-reacting with the S-glycoprotein of at least eight β-coronaviruses from lineages A, B and C, including all five human-infecting β-coronaviruses. This mAb, designated B6, broadly inhibits entry of viral particles pseudotyped with the S glycoprotein of lineage C (MERS-CoV and HKU4) and lineage A (OC43) coronaviruses, providing proof-of-concept of mAb-mediated broad β-coronavirus neutralization. A cryoEM structure of MERS-CoV S bound to B6 reveals that the mAb recognizes a linear epitope in the stem helix within a highly dynamic region of the S_2_ fusion machinery. Crystal structures of B6 in complex with MERS-CoV S, SARS-CoV/SARS-CoV-2 S, OC43 S and HKU4 S stem helix peptides combined with binding assays reveal an unexpected binding mode to a cryptic epitope, delineate the molecular basis of cross-reactivity and rationalize observed binding affinities for distinct coronaviruses. Collectively, our data indicate that B6 sterically interferes with S conformational changes leading to membrane fusion and identify a key target for next-generation structure-guided design of a pan-coronavirus vaccine.

## Results

### Isolation of a broadly neutralizing coronavirus mAb

To elicit cross-reactive Abs targeting conserved coronavirus S epitopes, we immunized mice twice with the prefusion-stabilized MERS-CoV S ectodomain trimer and once with the prefusion-stabilized SARS-CoV S ectodomain trimer **(Figure 1A)**. We subsequently generated hybridomas from immunized animals and implemented a selection strategy to identify those secreting Abs recognizing both MERS-CoV S and SARS-CoV S but not their respective S_1_ subunits (which are much less conserved than the S_2_ subunit(Walls et al., 2020b; Walls et al., 2016a)), the shared foldon trimerization domain or the his tag. We identified and sequenced a mAb, designated B6, that bound prefusion MERS-CoV S (lineage C) and SARS-CoV S (lineage B) trimers, the two immunogens used, as well as SARS-CoV-2 S (lineage B) and OC43 S (lineage A) trimers with nanomolar to picomolar avidities. Specifically, B6 bound most tightly to MERS-CoV S **(Figure 1B)**, followed by OC43 S (with one order of magnitude lower apparent affinity, **Figure 1C**) and SARS-CoV/SARS-CoV-2 S (with three orders of magnitude reduced apparent affinity, **Figure 1D-E**). These results show that B6 is a broadly reactive mAb recognizing at least four distinct S glycoproteins distributed across three lineages of the β-coronavirus genus.

**Figure 1.**
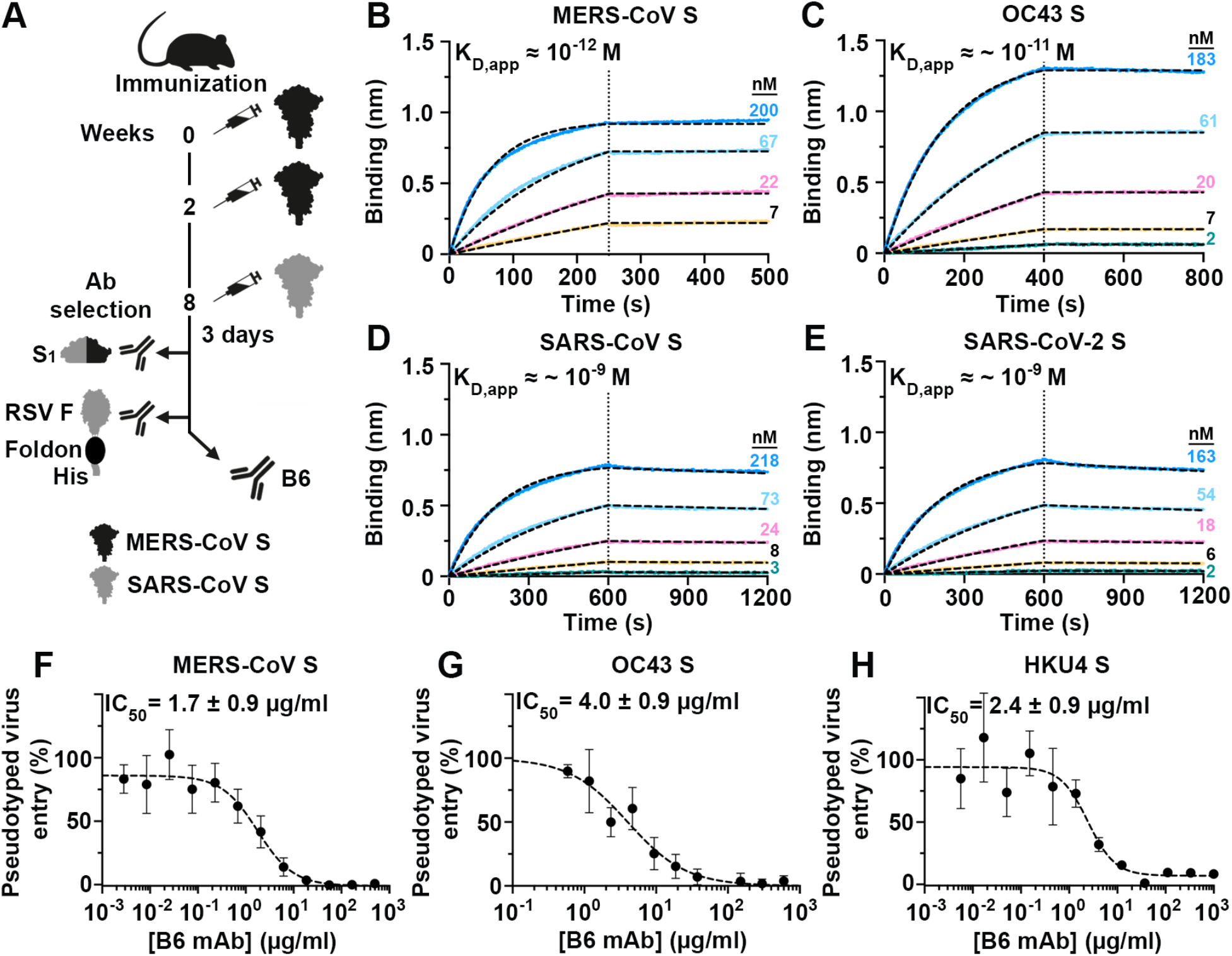
Identification and characterization of a cross-reactive and broadly neutralizing coronavirus mAb. (**A**) Mouse immunization and B6 mAb selection scheme. MERS-CoV and SARS-CoV S_1_ subunits fused to human Fc and the respiratory syncytial virus fusion glycoprotein (RSV F) ectodomain trimer fused to a foldon and a his-tag were used as decoys during selection. (**B-E**) Binding of MERS-CoV S (**B**), OC43 S (**C**), SARS-CoV S (**D**) and SARS-CoV-2 S (**E**) ectodomain trimers to the B6 mAb immobilized at the surface of biolayer interferometry biosensors. Data were analyzed with the ForteBio software, and global fits are shown as dashed lines. The vertical dotted lines correspond to the transition between the association and dissociation phases. Approximate apparent equilibrium dissociation constants (K_D_, app) are reported due to the binding avidity resulting from the trimeric nature of S glycoproteins. (**F-H)** B6-mediated neutralization of VSV particles pseudotyped with MERS-CoV S (**F**), OC43 S (**G**) and HKU4 S (**H**). Data were evaluated using a non-linear sigmoidal regression model with variable Hill slope. Fit is shown as dashed lines and experiments were performed in triplicate with at least two independent mAb and pseudotyped virus preparations.

To evaluate the neutralization potency and breadth of B6, we assessed S-mediated entry into cells of either vesicular stomatitis virus (VSV) (Kaname et al., 2010) or murine leukemia virus (MLV) (Millet and Whittaker, 2016; Walls et al., 2020b) pseudotyped with MERS-CoV S, OC43 S, SARS-CoV S, SARS-CoV-2 S and HKU4 S in the presence of varying concentrations of mAb. We determined half-maximal inhibitory concentrations of 1.7 ± 0.9 µg/mL, 4.0 ± 0.9 µg/mL and 2.4 ± 0.9 µg/mL for MERS-CoV S, OC43 S and HKU4 S pseudotyped viruses, respectively **(Figure 1F-G)** whereas no neutralization was observed for SARS-CoV S and SARS-CoV-2 S **(Figure S1)**. B6 therefore broadly neutralizes S-mediated entry of pseudotyped viruses harboring β-coronavirus S glycoproteins from lineages A and C, but not from lineage B, putatively due to lower-affinity binding **(Figure 1B-E)**.

### B6 targets a linear epitope in the fusion machinery

To identify the epitope recognized by B6, we determined a cryo-EM structure of the MERS-CoV S glycoprotein in complex with the B6 Fab fragment at 2.5 Å overall resolution (**Figure 2A-B, Figure S2 and Table 1**). 3D classification of the cryoEM data revealed incomplete Fab saturation, with one to three B6 Fabs bound to the MERS-CoV S trimer, and a marked conformational dynamic of bound B6 Fabs, yielding a continuum of conformations. Although these two factors compounded local resolution of the S/B6 interface, we identified that the B6 epitope resides in the stem helix (i.e. downstream from the connector domain and before the heptad-repeat 2 region) within the S_2_ subunit (so-called fusion machinery) **(Figure 2A-B)**. Our 3D reconstructions further suggest that B6 binding disrupts the stem helix quaternary structure, which is presumed to form a 3-helix bundle (observed in the NL63 S(Walls et al., 2016b) and SARS-CoV/SARS-CoV-2 S structures(Gui et al., 2017; Kirchdoerfer et al., 2018; Walls et al., 2020b; Walls et al., 2019; Wrapp et al., 2020; Yuan et al., 2017)) but not maintained in the B6-bound MERS-CoV S structure **(Figure 2A)**.

**Table 1.**
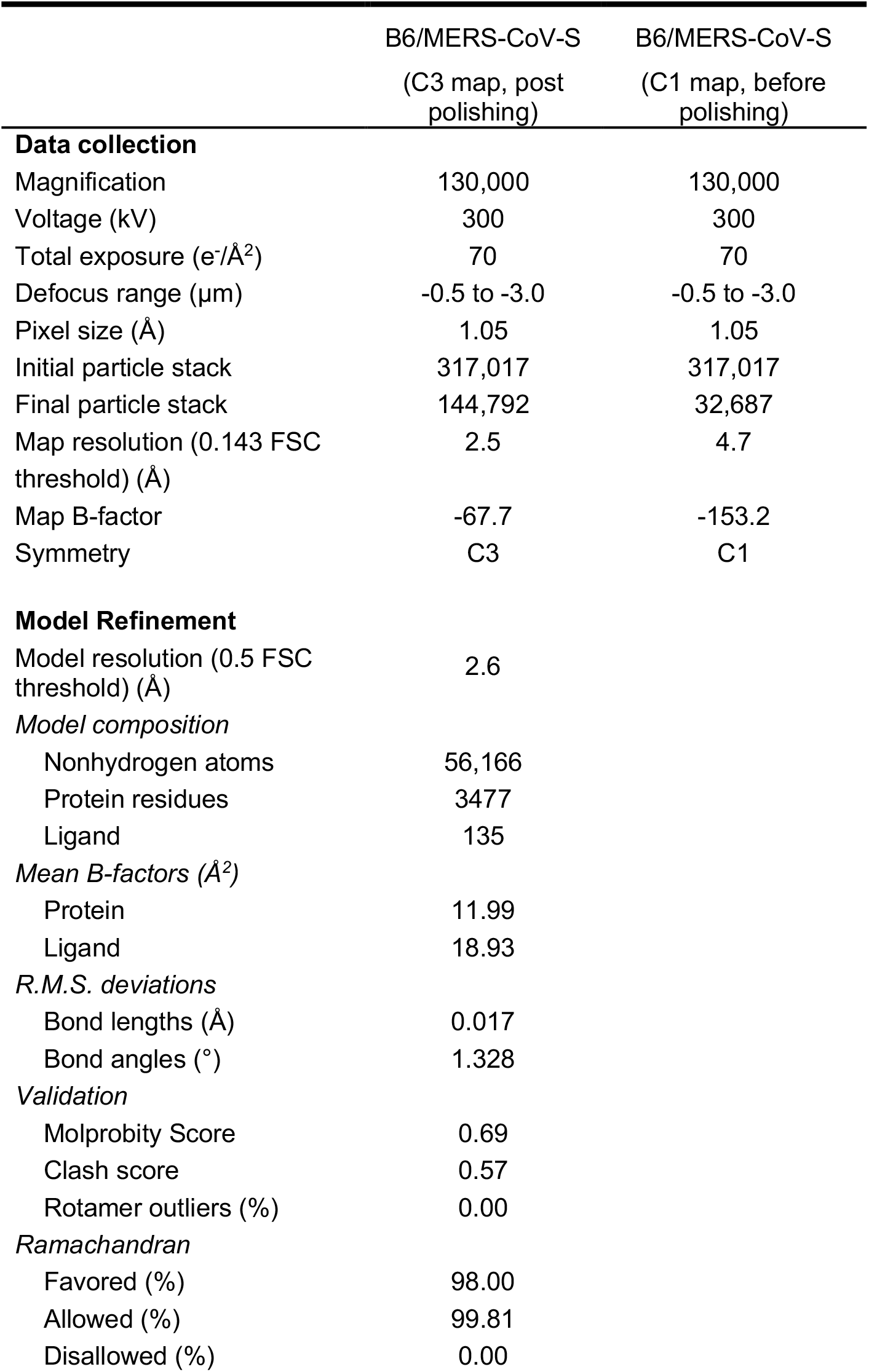
CryoEM data collection and refinement statistics.

**Figure 2.**
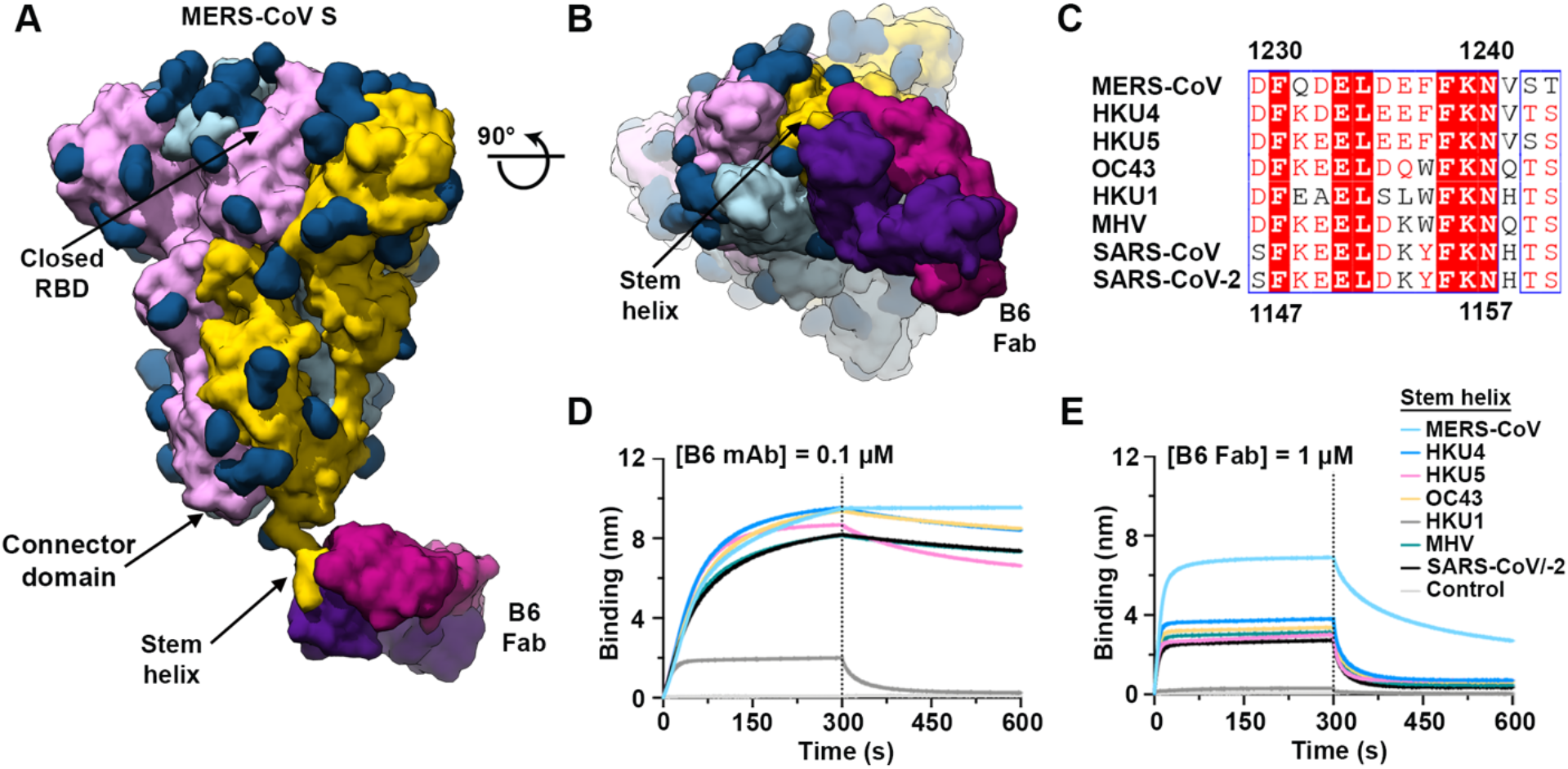
B6 targets a linear epitope in the coronavirus S_2_ fusion machinery. (**A-B**) Molecular surface representation of a composite model of the B6-bound MERS-CoV S cryoEM structure and of the B6-bound MERS-CoV S stem helix peptide crystal structure shown from the side (**A**) and viewed from the viral membrane (**B**). MERS-CoV S protomers are colored pink, cyan and gold and the B6 Fab heavy and light chains are colored purple and magenta, respectively. The composite model was generated by docking the crystal structure of B6 bound to the MERS-CoV stem helix in the cryoEM map. (**C**) Identification of a conserved 15 residue sequence spanning the stem helix. Residue numbering for MERS-CoV S and SARS-CoV-2 S are indicated on top and bottom of the alignment, respectively. (**D**) Binding of 0.1 µM B6 mAb or (**E**) 1 µM B6 Fab to biotinylated coronavirus S stem helix peptides immobilized at the surface of biolayer interferometry biosensors.

Based on our cryoEM structure, we identified a conserved 15 residue sequence at the C-terminus of the last residue resolved in previously reported MERS-CoV S structures(Pallesen et al., 2017; Park et al., 2019; Walls et al., 2019; Yuan et al., 2017) and confirmed by biolayer interferometry that it encompasses the B6 epitope using synthetic MERS-CoV S biotinylated peptides **(Figure 2C-E and Figure S3)**. We further found that B6 bound to the corresponding stem helix peptides from all known human-infecting β-coronaviruses: SARS-CoV-2 and SARS-CoV, the sequence is strictly conserved among the two viruses, OC43 and HKU1 as well as mouse hepatitis virus and two MERS-CoV-related bat viruses (HKU4 and HKU5) in mAb and Fab formats **(Figure 2D-E)**. B6 interacted most efficiently with the MERS-CoV S peptide, likely due to its major role in elicitation of this mAb, followed by all other coronavirus peptides tested, which bound with comparable affinities, except for HKU1 which interacted more weakly than other stem helix peptides.

### B6 recognizes a conserved epitope in the stem helix

To obtain an atomic-level understanding of the broad B6 cross-reactivity, we determined five crystal structures of the B6 Fab in complex with peptide epitopes derived from MERS-CoV S (residues 1230-1240 or 1230-1244), SARS-CoV S (residues 1129-1143), SARS-CoV-2 S (residues 1147-1161), OC43 S (residues 1232-1246) and HKU4 S (residues 1231-1245), at resolutions ranging from 1.4 to 1.8 Å **(Figure 3 A-F, Figure S4 and Table 2)**. In all five structures, the stem helix epitope folds as an amphipatic α-helix resolved for residues 1230-1240 (MERS-CoV S numbering) irrespective of the peptide length used for co-crystallization. B6 interacts with the helical epitope through shape complementarity, hydrogen-bonding and salt bridges using complementarity determining regions CDRH1-H3, framework region 3, CDRL1 and CDRL3 to bury ∼600Å^2^ at the paratope/epitope interface. The stem helix docks its hydrophobic face, lined by residues F1231_MERS-CoV_, L1235_MERS-CoV_, F1238_MERS-CoV_ and F1239_MERS-CoV_, into a hydrophobic groove formed by B6 heavy chain residues Y35, W49, V52 and L61 as well as light chain Y103 (**Figure 2C and 3A, B and D**). Moreover, B6 binding leads to the formation of a salt bridge triad, involving residue D1236_MERS-CoV_, CDRH3 residue R104 and CDRL1 residue H33.

**Table 2.** X-ray crystallography data collection and refinement statistics.

| Complex | B6/MERS-CoV 11aa | B6/MERS-CoV 15aa | B6/HKU4 | B6/OC43 | B6/SARS-CoV/SARS-CoV-2 |
| --- | --- | --- | --- | --- | --- |
| **Data collection** |  |  |  |  |  |
| Space group | C 1 2 1 | C 1 2 1 | C 1 2 1 | C 1 2 1 | C 1 2 1 |
| Cell constants<br>a,b,c (Å) | 95.06, 60.99,<br>80.26 | 93.591, 60.444,<br>79.71 | 93.59, 60.6,<br>79.77 | 92.99, 60.49,<br>79.39 | 93.18, 60.36,<br>79.70 |
| $\alpha, \beta, \gamma$ (°) | 90, 93.62,<br>90 | 90, 93.748,<br>90 | 90, 93.80,<br>90 | 90, 94.75,<br>90 | 90, 93.63,<br>90 |
| Wavelength (Å) | 1.000030 | 0.977410 | 0.977410 | 0.977410 | 0.977410 |
| Resolution (Å) | 43.89 -1.55<br>(1.61-1.55) | 43.49-1.35<br>(1.4-1.35) | 43.56-1.5<br>(1.55-1.5) | 43.57-1.8<br>(1.86-1.8) | 46.5-1.4<br>(1.45-1.4) |
| Rmerge (%) | 5.336 (49.25) | 3.514 (62.63) | 3.013 (49.45) | 4.765 (44.28) | 1.843 (39.76) |
| I/$\sigma$(I) | 6.78 (1.34) | 18.12 (1.44) | 9.13 (1.08) | 8.25 (1.20) | 12.70 (1.28) |
| CC(1/2) | 0.994 (0.64) | 1 (0.547) | 0.999 (0.607) | 0.998 (0.624) | 1 (0.816) |
| Completeness (%) | 99.91 (99.97) | 98.90 (93.96) | 96.77 (94.19) | 99.07 (97.35) | 98.61 (95.06) |
| Redundancy | 2.0 (2.0) | 3.2 (2.5) | 1.9 (1.9) | 1.9 (1.9) | 1.9 (1.9) |
| **Refinement** |  |  |  |  |  |
| Resolution (Å) | 43.89 -1.55 | 43.49-1.35 | 43.56-1.5 | 43.57-1.8 | 46.5-1.4 |
| Unique reflections | 66,514 | 95,828 | 69,063 | 40,478 | 85,691 |
| Rwork/Rfree (%) | 14.49/19.28 | 17.73/20.60 | 16.25/19.34 | 17.35/22.74 | 14.05/17.36 |
| Number of protein atoms | 3580 | 3574 | 3601 | 3562 | 3631 |
| Number of waters | 514 | 554 | 513 | 555 | 534 |
| R.m.s.d. bond lengths (Å) | 0.015 | 0.013 | 0.016 | 0.006 | 0.013 |
| R.m.s.d. bond angles (°) | 1.29 | 1.31 | 1.48 | 0.88 | 1.35 |
| Ramachandran favored (%) | 97.75 | 98.64 | 98.65 | 97.5 | 98.19 |
| Ramachandran allowed (%) | 2.25 | 1.36 | 1.35 | 2.5 | 1.81 |
| Ramachandran outliers (%) | 0 | 0 | 0 | 0 | 0 |

<sup>a</sup>Numbers in parentheses refer to outer resolution shell

**Figure 3.**
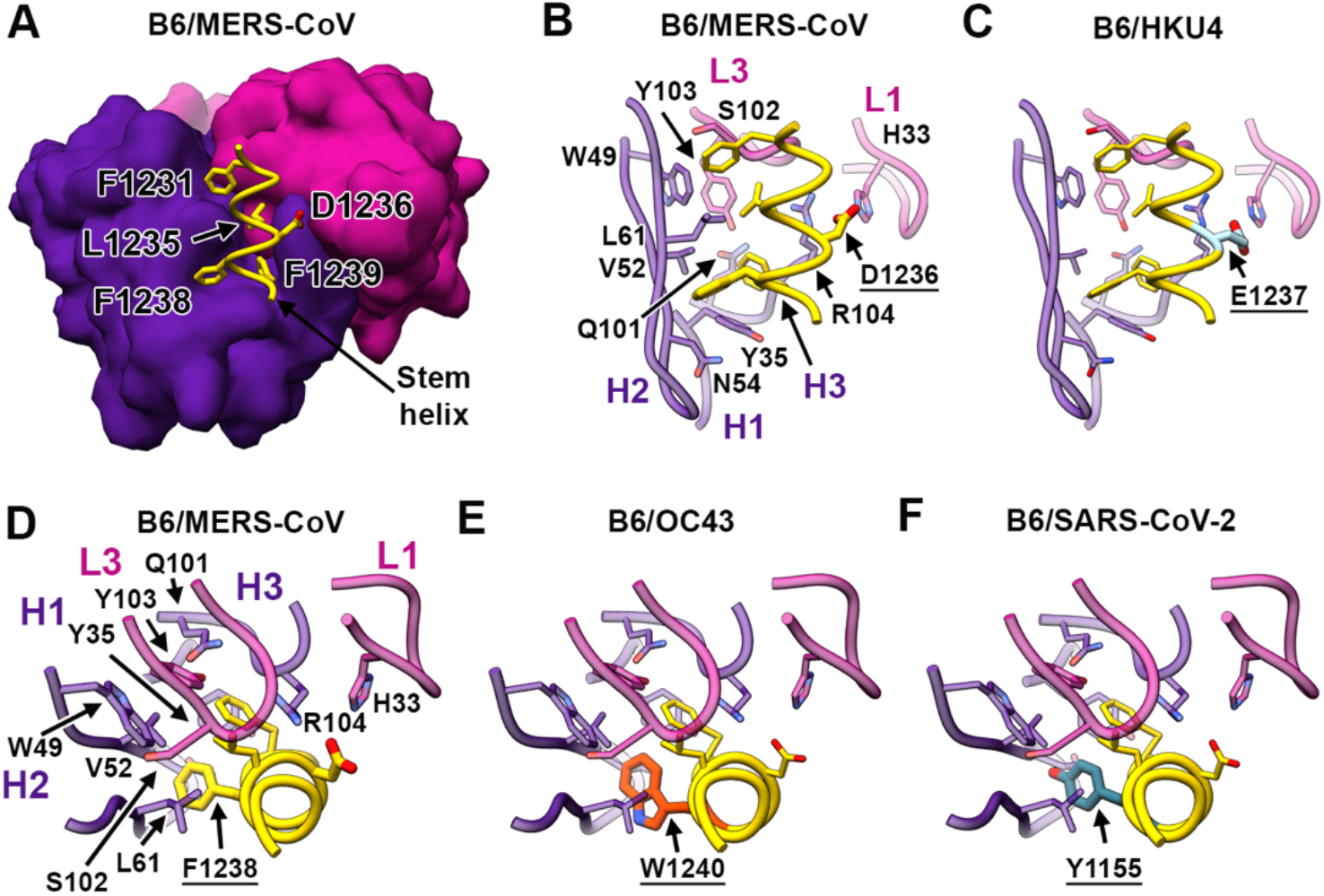
Molecular basis for the broad B6 cross-reactivity with a conserved coronavirus stem helix peptide. (**A**) Crystal structure of the B6 Fab (surface rendering) in complex with the MERS-CoV S stem helix peptide. (**B-C**) Crystal structures of the B6 Fab bound to the MERS-CoV S (B) or HKU4 S (C) stem helix reveal a conserved network of interactions except for the substitution of D1236_MERS-CoV_ with E1237_HKU4_ which preserves the salt bridge triad formed with CDRH3 residue R104 and CDRL1 residue H33. (**D-F**) Crystal structures of the B6 Fab bound to the MERS-CoV S (D), OC43 S (E) or SARS-CoV/SARS-CoV-2 S (F) stem helix showcasing the conservation of the paratope/epitope interface except for the conservative substitution of F1238_MERS-CoV_ with W1240_OC43_ or Y1137_SARS-CoV_/Y1155_SARS-CoV-2_. The B6 heavy and light chains are colored purple and magenta, respectively, and only selected regions are shown in panels (B-F) for clarity. The coronavirus S stem helix peptides are rendered in ribbon representation and colored gold with interacting side chains shown in stick representation.

Comparison of the B6-bound structures of MERS-CoV, HKU4, SARS-CoV/SARS-CoV-2 and OC43 S stem helix peptides explains the broad mAb cross-reactivity with β-coronavirus S glycoproteins as shape complementarity is maintained through strict conservation of 3 out of 4 hydrophobic residues whereas F1238_MERS-CoV_ is conservatively substituted with Y1137_SARS-CoV_/Y1155_SARS-CoV-2_ or W1240 _OC43_/W1237_HKU1_ (our structures demonstrate that all three aromatic side chains are accommodated by B6). Furthermore, the D1236_MERS-CoV_-mediated salt bridge triad is preserved, including with a non-optimal E1237_HKU4_ side chain, with the exception of S1235_HKU1_ which abrogates these interactions and explains the dampened B6 binding to the HKU1 peptide **(Figure 2C-E and 3 B-F)**.

B6 heavy chain residue L61 and CDRL1 residue H33 are mutated from germline and make major contributions to epitope recognition, highlighting the key contribution of affinity maturation to the cross-reactivity of this mAb.

### Mechanism of B6-mediated neutralization

We set out to elucidate the molecular basis of the B6-mediated broad neutralization of multiple coronaviruses from lineages A and C and lack of inhibition of lineage B coronaviruses. Our biolayer interferometry data indicate that although the B6 mAb efficiently interacted with the stem helix peptide of all but one of coronaviruses evaluated (HKU1, **Figure 2D-E**), the SARS-CoV-2 S and SARS-CoV S ectodomain trimers bound to B6 with three orders of magnitude reduced avidities compared to MERS-CoV S **(Figure 1B-E)**. Whereas the B6 epitope is not resolved in any prefusion coronavirus S structures determined to date, the stem helix region directly upstream is resolved to a much greater extent for SARS-CoV-2 S and SARS-CoV S, indicating a rigid structure(Gui et al., 2017; Kirchdoerfer et al., 2018; Walls et al., 2020b; Walls et al., 2019; Yuan et al., 2017) compared to MERS-CoV S (Pallesen et al., 2017; Park et al., 2019; Walls et al., 2019; Yuan et al., 2017), OC43 S (Tortorici et al., 2019), HKU1 S (Kirchdoerfer et al., 2016) or MHV S (Walls et al., 2016a) (**Figure 4A-C**). Furthermore, we determined B6 Fab binding affinities of 0.3 µM and 1.5 µM for MERS-CoV S and OC43 S, respectively, whereas SARS-CoV S recognition was too weak to accurately quantitate **(Figure S5)**. These findings along with the largely hydrophobic nature of the B6 epitope, which is expected to be occluded in the center of a 3-helix bundle **(Figure 4 D-E)** (as is the case for the region directly N-terminal to it), suggest that B6 recognizes a cryptic epitope and that binding to S trimers is modulated (at least in part) by the quaternary structure of the stem. The reduced conformational dynamics of the SARS-CoV-2 S and SARS-CoV S stem helix quaternary structure is expected to limit B6 accessibility to its cryptic epitope relative to other coronavirus S glycoproteins (**Figure 4A-E**). This hypothesis is supported by the correlation between neutralization potency and binding affinity which likely explains the lack of neutralization of lineage B β-coronaviruses.

**Figure 4.**
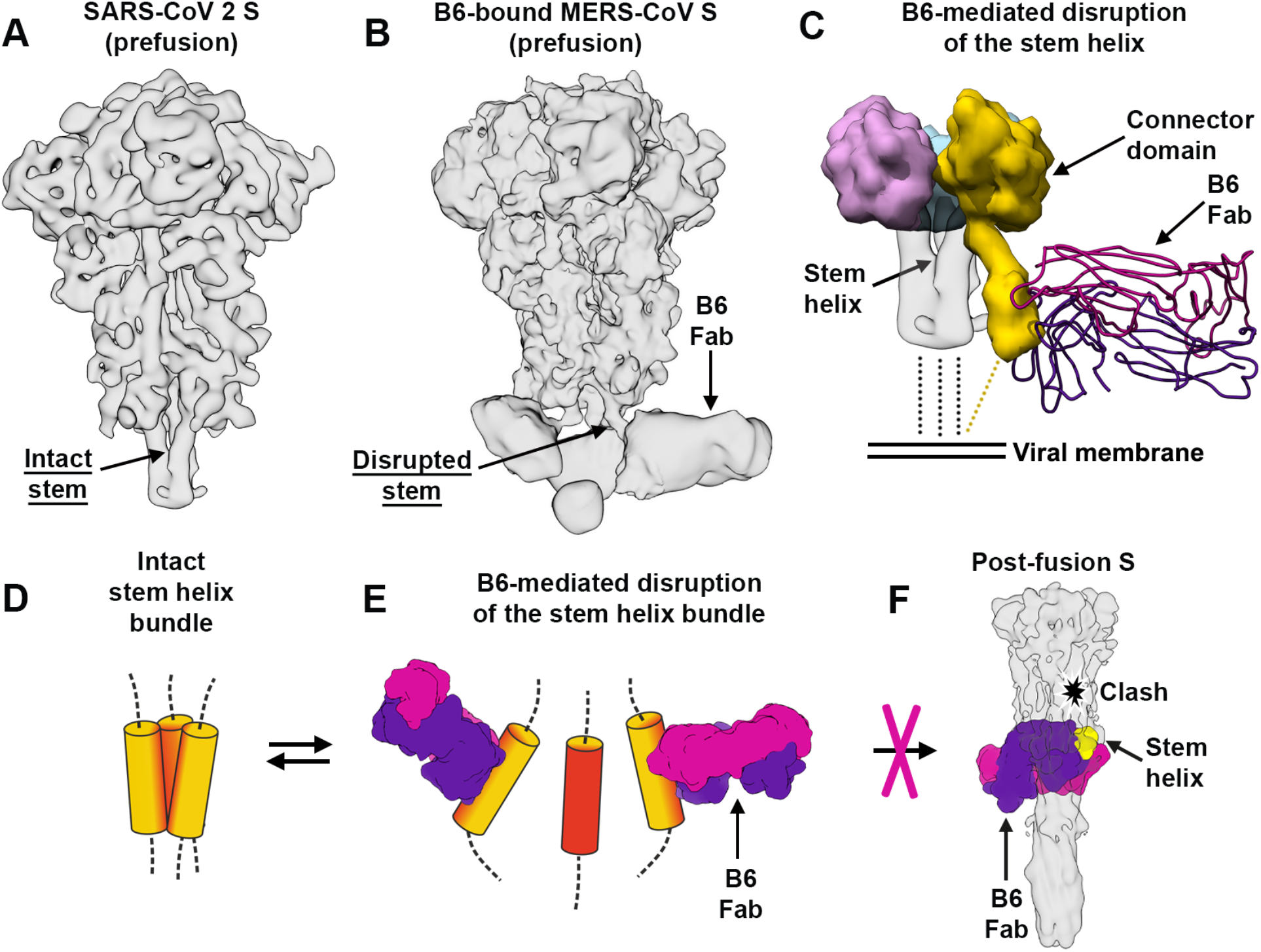
B6 binding disrupts the stem helix bundle and sterically inhibits membrane fusion. (**A**) CryoEM map of prefusion SARS-CoV-2 S (EMD-21452) filtered at 6 Å resolution to emphasize the intact trimeric stem helix bundle. (**B**) CryoEM map of the MERS-CoV S–B6 complex showing a disrupted stem helix bundle. (**C**) Model of B6-induced S stem movement obtained through comparison of the apo SARS-CoV-2 S and B6-bound MERS-CoV S structures. (**D-F**) Proposed mechanism of inhibition mediated by the B6 mAb. B6 binds to the hydrophobic core (red) of the stem helix bundle and disrupts its quaternary structure (D-E). The B6 disrupted state likely prevents S_2_ subunit refolding from the pre- to the post-fusion state and blocks viral entry (F).

Analysis of the postfusion mouse hepatitis S (Walls et al., 2017), SARS-CoV-2 S (Cai et al., 2020) and SARS-CoV S (Fan et al., 2020) structures show that the B6 epitope is buried at the interface with the other two protomers of the rod-shaped trimer. As a result, B6 binding appears to be incompatible with adoption of the postfusion S conformation (**Figure 4F**). Collectively, the data presented here suggest that B6 binding sterically interferes with S fusogenic conformational changes and likely block viral entry through inhibition of membrane fusion (**Figure 4C-F**), as proposed for fusion machinery-directed mAbs against influenza virus(Corti et al., 2011), ebolavirus(King et al., 2019) or HIV(Kong et al., 2016).

## Discussion

The high sequence variability of viral glycoproteins was long considered as an unsurmountable obstacle to the development of mAb therapies or vaccines conferring broad protection(Corti and Lanzavecchia, 2013). The identification of broadly neutralizing mAbs targeting conserved HIV-1 envelope epitopes from infected individuals brought about a paradigm shift for this virus undergoing extreme antigenic drift(Huang et al., 2012; Kong et al., 2016; Scheid et al., 2009; Walker et al., 2011; Walker et al., 2009; Wu et al., 2010; Zhou et al., 2010). Heterotypic influenza virus neutralization was also described for human cross-reactive mAbs recognizing the hemagglutinin receptor-binding site or the fusion machinery(Corti et al., 2011; Dreyfus et al., 2012; Ekiert et al., 2011; Ekiert et al., 2012; Kallewaard et al., 2016; Whittle et al., 2011). These findings were paralleled by efforts to identify broadly neutralizing Abs against respiroviruses(Corti et al., 2013), henipaviruses(Dang et al., 2019; Mire et al., 2019; Zhu et al., 2006), Dengue and Zika viruses(Barba-Spaeth et al., 2016; Dejnirattisai et al., 2015; Rouvinski et al., 2015) or ebolaviruses(Bornholdt et al., 2016; Flyak et al., 2018; King et al., 2019; West et al., 2018).

The genetic diversity of coronaviruses circulating in chiropteran and avian reservoirs along with the recent emergence of multiple highly pathogenic coronaviruses showcase the need for vaccines and therapeutics that protect humans against a broad range of viruses. As the S_2_ fusion machinery contains several important antigenic sites and is more conserved than the S_1_ subunit, it is an attractive target for broad-coronavirus neutralization(Tortorici and Veesler, 2019; Walls et al., 2016a). Previous studies described conserved epitopes targeted by neutralizing Abs, such as the fusion peptide or heptad-repeats, as well as a variable loop in the MERS-CoV S connector domain (Daniel et al., 1993; Elshabrawy et al., 2012; Pallesen et al., 2017; Poh et al., 2020; Walls et al., 2016a; Wec et al., 2020; Zhang et al., 2004; Zheng et al., 2020). The discovery of the B6 mAb provides proof-of-concept of mAb-mediated broad β-coronavirus neutralization and uncovers a previously unknown conserved cryptic epitope that is predicted to be located in the hydrophobic core of the stem helix. B6 cross-reacts with at least eight distinct S glycoproteins, from β-coronaviruses belonging to lineages A, B and C, and broadly neutralize two human and one bat pseudotyped viruses from lineages A and C. B6 could be used for detection or diagnostic of coronavirus infection and humanized versions of this mAb are promising candidate therapeutics against emerging and re-emerging β-coronaviruses from lineages A and C. Our data further suggest that affinity maturation of B6 using SARS-CoV-2 S and SARS-CoV S might enhance recognition of and extend neutralization breadth towards β-coronaviruses from lineage B. Finally, the identification of the conserved B6 epitope paves the way for epitope-focused vaccine design(Azoitei et al., 2011; Correia et al., 2014; Sesterhenn et al., 2020) that could elicit pan-coronavirus immunity, as supported by the elicitation of the B6 mAb through vaccination and the recent findings that humans and camels infected with MERS-CoV, humans infected with SARS-CoV-2 and humanized mice immunized with a cocktail of coronavirus S glycoproteins produce antibodies targeting an epitope similar to the one targeted by B6(Song et al., 2020; Wang et al., 2020b).

## Acknowledgments

We thank Hideki Tani (University of Toyama) for providing the reagents necessary for preparing VSV pseudotyped viruses and Brooke Fiala for assisting with protein production. This study was supported by the National Institute of General Medical Sciences (R01GM120553 to D.V.), the National Institute of Allergy and Infectious Diseases (DP1AI158186 and HHSN272201700059C to D.V.), a Pew Biomedical Scholars Award (D.V.), an Investigators in the Pathogenesis of Infectious Disease Awards from the Burroughs Wellcome Fund (D.V.), a Fast Grants (D.V.), the University of Washington Arnold and Mabel Beckman cryoEM center, the Swiss National Science Foundation (P400PB_183942 to M.M.S.), the Pasteur Institute (M.A.T.) the M.J. Murdock Charitable Trust (A.T.M and B.H.), and beamlines 8.2.1 and 5.0.1 at the Advanced Light Source at Lawrence Berkley National Laboratory.

## Supplementary Information

**Figure S1.**
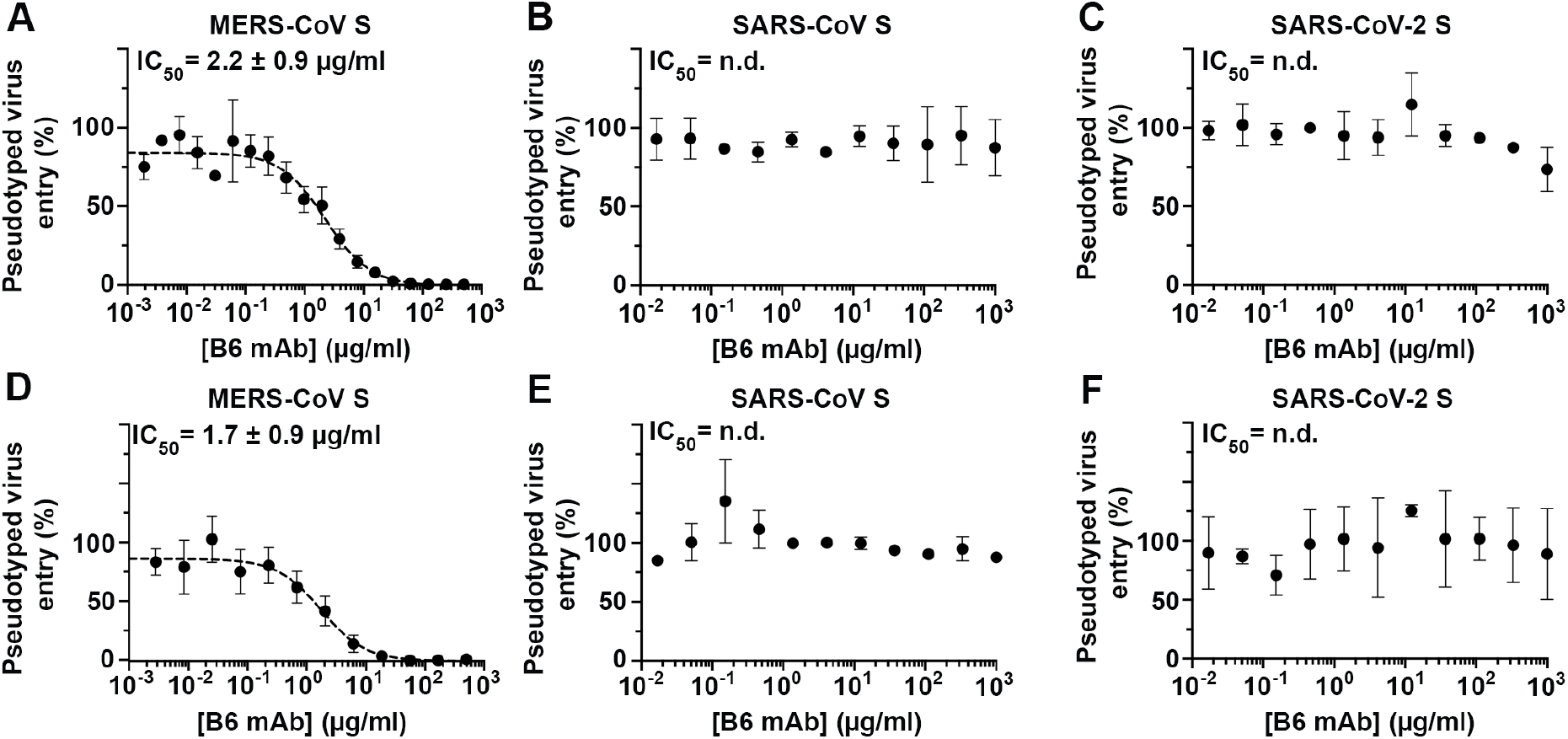
MERS-CoV S, SARS-CoV S and SARS-CoV-2 S pseudotyped virus neutralization. Neutralization assays of MLV (**A-C**) or VSV (**D-F**) particles pseudotyped with (**A**,**D**) MERS-CoV S (**B**,**E**) SARS-CoV S and (**C**,**F**) SARS-CoV-2 S were performed in the presence of the indicated concentration of B6 mAb. Data were evaluated using a non-linear sigmoidal regression model with variable Hill slope. Experiments were performed in triplicates with at least two independent mAb and pseudotyped virus preparations.

**Figure S2.**
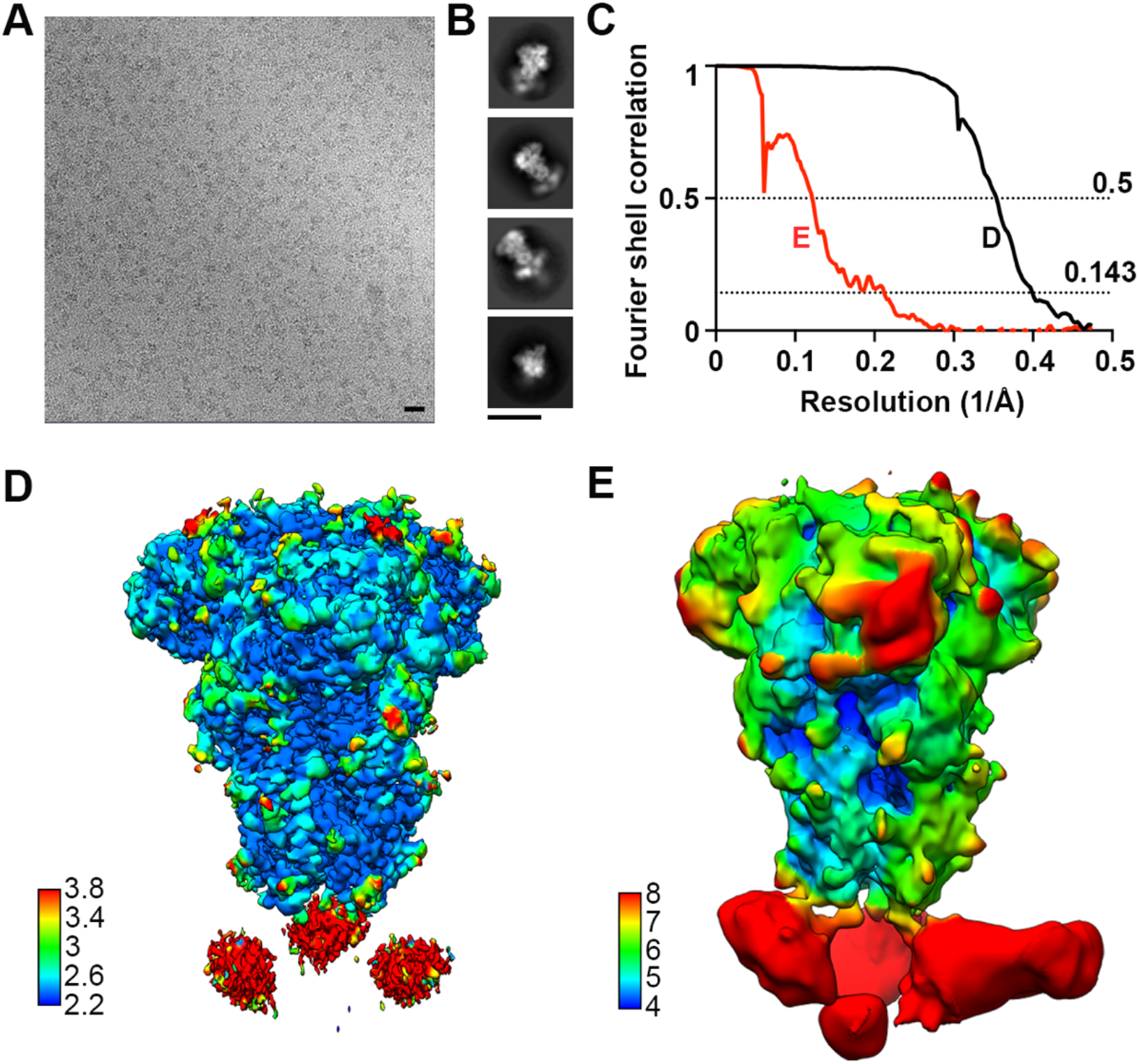
CryoEM characterization of the B6-bound MERS-CoV S complex. (**A**) Representative cryoEM micrograph of the MERS-CoV S prefusion trimer bound to B6 embedded in vitreous ice. Scale bar: 20 nm. (**B**) Selected reference-free 2D class averages. Scale bar: 20 nm. (**C**) Fourier shell correlation curves for the reconstructions shown in panels D and E. **(D)** Reconstruction obtained with all selected particles and applying C3 symmetry colored by local resolution. **(E)** Reconstruction obtained with a subset of particles obtained through focused classification to improve B6 resolvability colored by local resolution.

**Figure S3.**
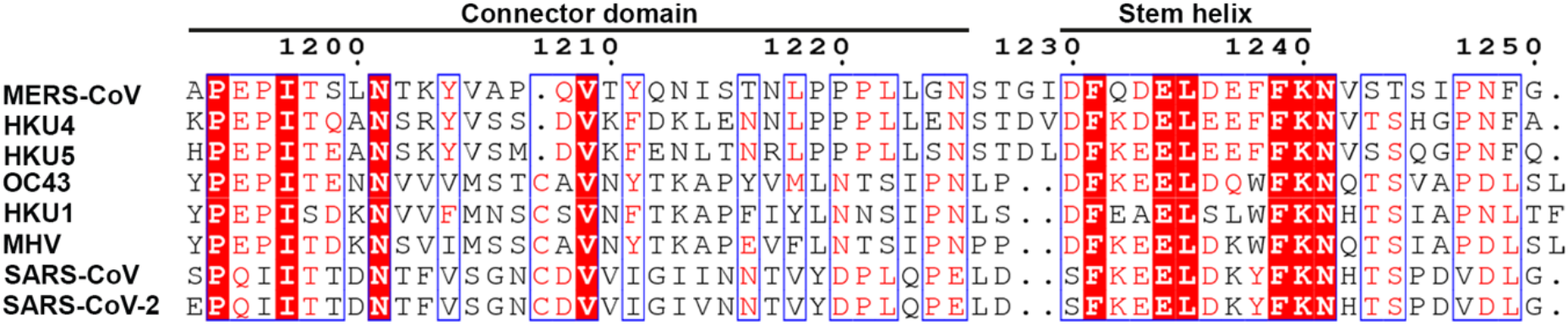
Protein sequence alignment of the stem region for selected β-coronavirus S glycoproteins. The sequence alignment was performed based on MERS-CoV S using the following S protein sequences: MERS-CoV EMC/2012 (GenBank: AFS88936.1), HKU4 (UniProtKB: A3EX94.1), HKU5 (UniProtKB: A3EXD0.1), HKU1 isolate N5 (UniProtKB: Q0ZME7.1), MHV A59 (UniProtKB: P11224.2), OC43 (UniProtKB: Q696P8), SARS-CoV Urbani (GenBank: AAP13441.1), SARS-CoV-2 (NCBI Reference Sequence: YP_009724390.1). Sequence alignment was performed using Multalin(Corpet, 1988) and visualized using ESPrint3.0(Robert and Gouet, 2014). The conserved stem helix recognized by B6 is indicated.

**Figure S4.**
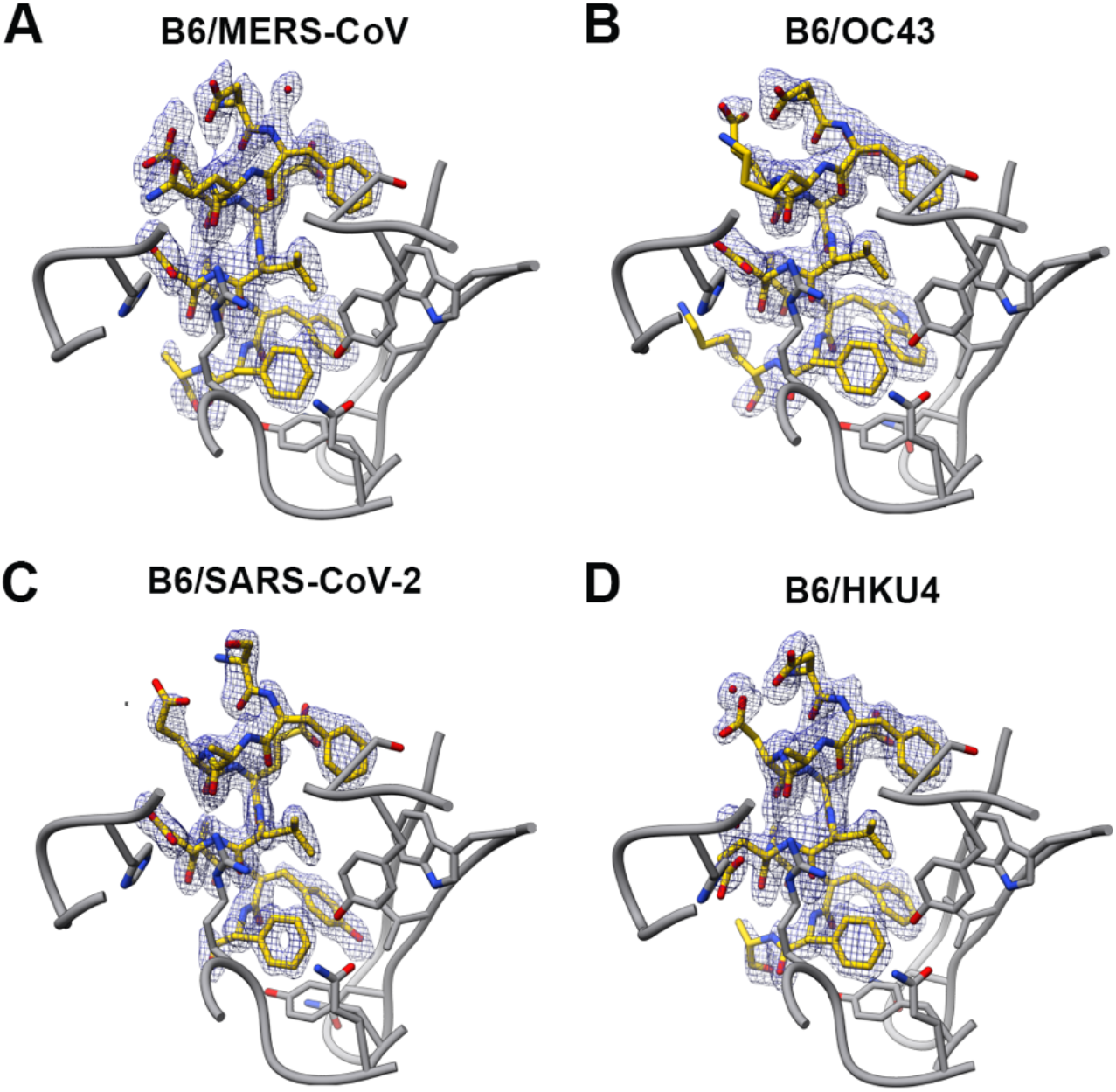
Crystal structures of B6 bound to coronavirus S stem helix peptides. Stem peptides of (**A**) MERS-CoV S (**B**) OC43 S (**C**) SARSCoV/SARS-CoV-2 S and (**D**) HKU4 S are shown in stick representation with carbon atoms colored yellow. B6 is shown in ribbon representation with interacting residues rendered as stick representation in gray. Oxygen and nitrogen atoms are colored red and blue, respectively. The 2Fo-Fc maps for the different peptides are shown as a blue mesh at a contour level of 1 σ.

**Figure S5.**
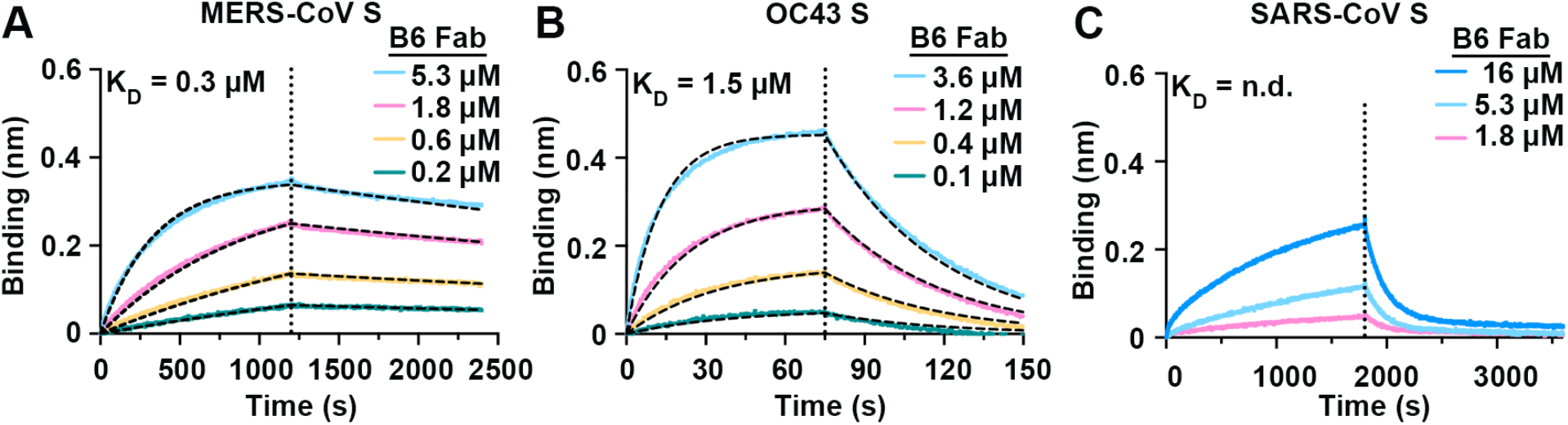
B6 binding kinetics to different coronavirus S ectodomain trimers. **A-C**) Binding of B6 to immobilized (**A**) MERS-CoV S, (**B**) OC43 S and (**C**) SARS-CoV S measured by biolayer interferometry. The vertical dotted lines correspond to the transition between the association and dissociation phases. Data are shown for one representative measurement and were analyzed with the OctetBio software. Global fits are shown as dashed lines. We determined dissociation constant (K_D_) values of 0.28 (0.2) ± 0.001 and 1.50 (1.47) ± 0.01 µM for two independent batches of S protein for MERS-CoV S and OC43 S, respectively. The dissociation constant for SARS-CoV S could not be evaluated reliably, however, the predicted affinity is significantly lower compared to the other two S proteins.

## Methods

### Identification of the B6 broadly neutralizing mAb

Ten-week-old CD-1 mice were injected twice with 50 µg of MERS-CoV S formulated with Adjuplex at weeks 0 and 2 and once with 50 µg of SARS-CoV S formulated with Adjuplex at week 8 at the Fred Hutchinson Cancer Research Center Antibody Technology Resource. 3 days after the final injection splenocytes were isolated from high titer mice and electrofused with P3×63-Ag8 myeloma cell line (BTX, Harvard Apparatus). Hybridoma supernatants were tested for binding to prefusion SARS-CoV S, MERS-CoV S, SARS-CoV S_1_ subunit, MERS-CoV S_1_ subunit and respiratory syncytial virus F (which harbors a foldon motif and a his tag similar to the SARS-CoV S and MERS-CoV S ectodomain trimer constructs) using a high throughput bead-based binding array. Hybridomas from wells containing supernatants that were positive for binding to prefusion SARS-CoV S and MERS-CoV S but negative for SARS-CoV S_1_, MERS-CoV S_1_, and respiratory syncytial virus F were sub-cloned by limiting dilution and re-screened for binding as above. The VH and VL sequences of B6 were recovered using the mouse iG primer set (Millipore) using the protocol outlined in (Siegel, 2009), and Sanger sequenced (Genewiz). The VH/VL sequences were codon-optimized and cloned into full-length pTT3 derived IgG1 and IgL kappa expression vectors containing human constant regions using Gibson assembly (Snijder et al., 2018).

### Protein expression and purification

MERS-CoV 2P S, OC43 S, SARS-CoV 2P S and SARS-CoV-2 2P S were produced as previously described (Tortorici et al., 2019; Walls et al., 2020b; Walls et al., 2019). Briefly, all ectodomains were produced in HEK293F cells grown in suspension using FreeStyle 293 expression medium (Life Technologies) at 37 °C in a humidified 8% (v/v) CO_2_ incubator rotating at 130 r.p.m. The cultures were transfected using 293fectin (ThermoFisher Scientific) with cells grown to a density of 10^6^ cells/ml and cultivated for three days. The supernatants were harvested and cells resuspended for another three days, yielding two harvests. For MERS-CoV 2P S, SARS-CoV 2P S and SARS-CoV-2 2P S, clarified supernatants were purified using a 5 ml Cobalt affinity column (Takara). HCoV-OC43 S was purified using a StrepTrap HP column (GE healthcare). Purified proteins were concentrated, flash-frozen in Tris-saline (50 mM Tris, pH 8.0 (25°C), 150 mM NaCl) and stored at −80°C. The MERS-CoV S_1_-Fc and SARS-CoV S_1_-Fc were previously described (Raj et al., 2013), produced as aforementioned for the prefusion S trimers and purified using protein A affinity chromatography.

For mAb B6 production, 250 µg of B6 heavy and 250 µg of B6 light chain encoding plasmids were co-transfected per liter of suspended HEK293F culture using 293Free transfection reagent (Millipore Sigma) according to the manufacturer’s instructions. Cells were transfected at a density of 10^6^ cells/ml. Expression was carried out for 6 days after which cells and cellular debris were removed by centrifugation at 4,000 × *g* followed by filtration through a 0.22 µm filter. Clarified cell supernatant containing recombinant mAb was passed over Protein A Agarose resin (Thermo Fisher Scientific). Protein A resin was extensively washed with 25 mM Phosphate pH 7.4, 150 mM NaCl (PBS) and eluted with IgG elution buffer (Thermo Scientific). Purified B6 was extensively dialyzed against PBS, concentrated, flash-frozen and stored at −80°C.

DS-Cav1-foldon-SpyTag (McLellan et al., 2013) was produced by lentiviral transduction of HEK293F cells using the Daedalus system (Bandaranayake et al., 2011). Lentivirus was produced by transient transfection of HEK293T cells (ATCC) using linear 25 kDa polyethyleneimine (PEI; Polysciences). Briefly, 4×10^6 cells were plated onto 10 cm tissue culture plates. After 24 h, 3 mg of psPAX2, 1.5 mg of pMD2G (Addgene plasmids #12260 and #12259, respectively), and 6 mg of lentiviral vector plasmid were mixed in 500 mL diluent (5 mM HEPES, 150 mM NaCl, pH 7.5) and 42 mL of PEI (1 mg/mL) and incubated for 15 minutes. The DNA/PEI complex was then added to the plate dropwise. Lentivirus was harvested 48 h post-transfection and concentrated 100× by centrifugation at 8000×g for 18 h. Transduction of the target cell line was carried out in 125 mL shake flasks containing 10×10^6 cells in 10 mL of growth media. 100 μL of 100× lentivirus was added to the flask and the cells were incubated with 225 rpm oscillation at 37°C in 8% CO_2_ for 4–6 hours, after which 20 mL of growth media was added to the shake flask. Transduced cells were expanded every other day to a density of 1×10^6 cells/mL until a final culture size of 4 L was reached. The media was harvested after 17 days of total incubation after measuring final cell concentration (∼5×10^6 cells/mL) and viability (∼90% viable). Culture supernatant was harvested by low-speed centrifugation to remove cells from the supernatant. NaCl and NaN3 were added to final concentrations of 250 mM and 0.02%, respectively. The supernatant was loaded over one 5 mL HisTrap FF Crude column (GE Healthcare) at 5 mL/min by an AKTA Pure (GE Healthcare). The 5 mL HisTrap column was washed with 10 column volumes of wash buffer (2× GIBCO 14200-075 PBS, 5 mM Imidazole, pH 7.5) followed by 6 column volumes of elution buffer (2× GIBCO 14200-075 PBS, 150 mM Imidazole, pH 7.5). The nickel elution was applied to a HiLoad 16/600 Superdex 200 pg column (GE Healthcare) and run in dPBS (GIBCO 14190-144) with 5% glycerol (Thermo BP229-1) to further purify the target protein by size-exclusion chromatography. The purified protein was snap frozen in liquid nitrogen and stored at −80°C.

### Kinetics of B6 mAb binding to coronavirus S proteins

The avidities of complex formation between B6 mAb and selected coronavirus S proteins were determined in PBS supplemented with 0.005 % Tween20 and 0.1% BSA (PBSTB) at 30 °C and 1,000 RPM shaking on an Octet Red instrument (Fortebio). Curve fitting was performed using a 1:1 binding model and the ForteBio data analysis software. K_D_ ranges were determined with a global fit. AHC biosensors (ForteBio) were hydrated in water and subsequently equilibrated in PBSTB buffer. 10 μg/mL B6 mAb was loaded to the biosensors to a shift of approximately 1nm. Then, the system was equilibrated in PBSTB buffer for 300 s prior to immersing the sensors in the respective coronavirus S protein (0 − 218 nM) for up to 600 s prior to dissociation in buffer for additional 600 s.

### Binding of B6 to different synthetic coronavirus S stem peptides

B6 binding analysis to selected biotinylated coronavirus S stem helix peptides was performed in PBS supplemented with 0.005 % Tween20 (PBST) at 30 °C and 1,000 RPM shaking on an Octet Red instrument (Fortebio).

1 µg/ml biotinylated stem peptide (15- or 16-residue long stem peptide-PEG6-Lys-Biotin synthesized fom Genscript) was loaded on SA biosensors to a threshold of 0.5 nm. Then, the system was equilibrated in PBST for 300 s prior to immersing the sensors in 0.1 µM B6 mAb or 1 µM B6 Fab, respectively, for 300 s prior to dissociation in buffer for 300 s.

### Kinetics of B6 Fab binding to different coronavirus S proteins

The rate constants of binding (*k*_on_) and dissociation (*k*_off_) for the complex between the B6 Fab and selected coronavirus S proteins were performed in PBST at 30 °C and 1,000 RPM shaking on an Octet Red instrument (Fortebio). Global curve fitting was performed using a 1:1 binding model and the ForteBio data analysis software. For MERS-CoV S and SARS-CoV S, HIS1K or Ni-NTA biosensors (ForteBio) were hydrated in water and subsequently equilibrated in PBST buffer. 20 μg/mL SARS-CoV S or 10 μg/mL MERS-CoV S, respectively, were loaded to the biosensors for up to 1800 s (1-4nm shift). The system was equilibrated in PBST for 300 s prior to immersing the sensors in B6 Fab (0 − 16 µM) for up to 1800 s prior to dissociation in buffer for 1800 s. For OC43 S, ARG2 biosensors were hydrated in water then activated for 300 s with an NHS-EDC solution (ForteBio) prior to amine coupling. 20 μg/mL OC43 was amine coupled to AR2G (ForteBio) sensors in 10 mM acetate pH 6.0 (ForteBio) respectively for 300 s and then quenched with 1M ethanolamine (ForteBio) for 300 s. The system was equilibrated in PBST for 300 s prior to immersing the sensors in B6 Fab (0 − 4 µM) for 75 s prior to dissociation in buffer for 75 s.

### Pseudovirus entry assays

Production of OC43 S pseudotyped VSV virus and the neutralization assay was performed as described previously (Hulswit et al., 2019; Tortorici et al., 2019). Briefly, HEK-293T cells at 70∼80% confluency were transfected with the pCAGGS expression vectors encoding full-length OC43 S with a truncation of the 17 C-terminal residues (to increase cell surface expression levels) along with fusion to a flag tag and the Fc-tagged bovine coronavirus hemagglutinin esterase protein at molar ratios of 8:1. 48 h after transfection, cells were transduced with VSVΔG/Fluc (bearing the *Photinus pyralis* firefly luciferase) (Kaname et al., 2010) at a multiplicity of infection of 1. Twenty-four hours later, supernatant was harvested and filtered through 0.45 μm membrane. Pseudotyped VSV virus was titrated on monolayer on HRT-18 cells. In the virus neutralization assay, serially diluted mAbs were pre-incubated with an equal volume of virus at room temperature for 1 h, and then inoculated on HRT-18 cells, and further incubated at 37°C. After 20 h, cells were washed once with PBS, lysed with cell lysis buffer (Promega) and firefly luciferase expression was measured on a Berthold Centro LB 960 plate luminometer using D-luciferin as a substrate (Promega). Percentage of infectivity was calculated as the ratio of luciferase readout in the presence of mAbs normalized to luciferase readout in the absence of mAb, and half maximal inhibitory concentrations (IC_50_) were determined using 4-parameter logistic regression (GraphPad Prism v8.0).

MERS-CoV S, SARS-CoV S and SARS-CoV-2 S pseudotyped VSV were prepared using 293T cells seeded in 10-cm dishes in DMEM supplemented with 10% FBS, 1% PenStrep and transfected with plasmids encoding for the corresponding S glycoprotein (24 µg/dish) using lipofectamine 2000 (Life Technologies) according to the manufacturer’s instructions. One day post-transfection, cells were infected with VSV(G*ΔG-luciferase). After 2 h, infected cells were washed four times with DMEM before medium supplemented with anti-VSV-G antibody (I1-mouse hybridoma supernatant diluted 1 to 50, from CRL-2700, ATCC). Particles were harvested 18 h post-inoculation, clarified from cellular debris by centrifugation at 2,000 x g for 5 min and used for neutralization experiments.

MERS-CoV S, SARS-CoV S, and SARS-CoV-2 S pseudotypes MLV were prepared as previously described (Walls et al., 2020b).

For viral neutralization, Huh7 cells (for MERS-CoV S pseudotyped virus) or stable 293T cells expressing ACE2 (Crawford et al., 2020) (for SARS-CoV S and SARS-CoV-2 S pseudotyped viruses) in DMEM supplemented with 10% FBS, 1% PenStrep were seeded at 40,000 cells/well into clear bottom white walled 96-well plates and cultured overnight at 37°C. Twelve-point 3-fold serial dilutions of B6 mAb were prepared in DMEM and pseudotyped VSV were added 1:1 to each B6 dilution in the presence of anti-VSV-G mAb from I1-mouse hybridoma supernatant diluted 50 times. After 45 min incubation at 37°C, 40 µl of the mixture was added to the cells and 2 h post-infection, 40 μL DMEM was added to the cells for 17-20 h. Following infection, medium was removed and 80 μL One-Glo-EX substrate (Promega) was added to the cells and incubated in the dark for 10 min prior reading on a Varioskan LUX plate reader (ThermoFisher).

### Western blots

SDS–PAGE (4x) loading buffer was added to all concentrated pseudovirus samples. The samples were run on a 4–20% (wt/vol) gradient Tris-glycine gel (BioRad) and transferred to PVDF membranes. B6 was used as primary Ab (1:500 dilution) and an Alexa Fluor 680-conjugated goat anti-human secondary Ab (1:50,000 dilution, Jackson Laboratory) were used for western blotting. A LI-COR processor was used to develop images.

### CryoEM sample preparation and data collection

Lacey carbon copper grids (400 mesh) were coated with a thin-layer of continuous carbon using a carbon evaporator. 1 mg/ml MERS-CoV S was incubated with 100 mM neuraminic acid (to promote the closed trimer conformation), 150mM Tris pH 8 (25°C) 150 mM NaCl for 16 h at 4°C. Then a 2-fold molar excess of B6 Fab over MERS-CoV S protomer was added to the solution and incubated for 1h at 37°C. The sample was diluted to 0.2 mg/ml S protein with 100 mM neuraminic acid-150mM Tris pH 8 (25°C) 150mM NaCl before 3 µl sample were applied on to a freshly glow discharged grid. Plunge freezing was performed using a TFS Vitrobot Mark IV (blot force: −1, blot time: 2.5 s, Humidity: 100 %, temperature: 25 °C). Data were acquired using an FEI Titan Krios transmission electron microscope operated at 300 kV and equipped with a Gatan K2 Summit direct detector and Gatan Quantum GIF energy filter, operated in zero-loss mode with a slit width of 20 eV. Automated data collection was carried out using Leginon (Suloway et al., 2005) at a nominal magnification of 130,000x with a pixel size of 0.525Å. The dose rate was adjusted to 8 counts/pixel/s, and each movie was acquired in super-resolution mode fractionated in 50 frames of 200 ms. 2,180 micrographs were collected in a single session with a defocus range comprised between −0.5 and −3.0 μm.

### CryoEM data processing

Movie frame alignment, estimation of the microscope contrast-transfer function parameters, particle picking and extraction were carried out using Warp (Tegunov and Cramer, 2019). Particle images were extracted with a box size of 800 pixels^2^ binned to 400 pixels^2^ yielding a pixel size of 1.05 Å. Two rounds of reference-free 2D classification were performed using Relion3.0 (Zivanov et al., 2018) to select well-defined particle images. Subsequently, two rounds of 3D classification with 50 iterations each (angular sampling 7.5° for 25 iterations and 1.8° with local search for 25 iterations), using the previously reported closed MERS-CoV S structure without the G4 Fab (PDB 5W9J) as initial model were carried out using Relion without imposing symmetry.

For the high resolution map, particle images were subjected to Bayesian polishing (Zivanov et al., 2019) before performing non-uniform refinement, defocus refinement and non-uniform refinement again in cryoSPARC (Punjani et al., 2017). Finally, two rounds of global CTF refinement of beam-tilt, trefoil and tetrafoil parameters was performed before a final round of non-uniform refinement to produce the 2.5Å resolution map.

For the lower resolution map, one additional round of focused classification in Relion with 50 iterations using a broad mask covering the region of interest (B6/stem) was carried out to further separate distinct B6 Fab conformations. 3D refinements of the best subclasses were carried out using homogenous refinement in cryoSPARC (Punjani et al., 2017). Reported resolutions are based on the gold-standard Fourier shell correlation (FSC) of 0.143 criterion and Fourier shell correlation curves were corrected for the effects of soft masking by high-resolution noise substitution (Chen et al., 2013).

### CryoEM model building and analysis

UCSF Chimera (Pettersen et al., 2004) and Coot (Emsley et al., 2010) were used to fit atomic models into the cryoEM maps. The MERS-CoV S EM structure in complex with 5-N-acetyl neuraminic acid (PDB 6Q04, residue 18-1224) and the B6-MERS-CoV_11_ (residue 1230-1240) crystal structure were fit into the cryoEM map. Subsequently the linker connecting the stem helix to the rest of the MERS-CoV S ectodomain (residue 1225-1229) was manually built using Coot. N-linked glycans were hand-built into the density where visible and the models were refined and relaxed using Rosetta using both sharpened and unsharpened maps (Frenz et al., 2019; Wang et al., 2016). Models were analyzed using MolProbity (Chen et al., 2010), EMringer (Barad et al., 2015), Phenix (Liebschner et al., 2019) and privateer (Agirre et al., 2015) to validate the stereochemistry of both the protein and glycan components. Figures were generated using UCSF Chimera.

### Crystallization and structure determination

All crystallization experiments were performed at 23 °C in hanging drop vapor diffusion experiments with initial concentrations of 20 mg/ml and 1.5-fold molar excess of peptide ligand. Crystal trays were setup with a mosquito using 100 nL mother liquor solution and 100 or 150 nL B6/peptide complex solution, respectively. Crystals of B6/MERS-CoV_11_ and B6/OC43_15_ appeared after several weeks in 0.2 M Potassium Thiocyanate and 20% (w/v) PEG3350, B6/MERS-CoV_15_ in 0.2 M Magnesium Chloride and 20% (w/v) PEG3350, B6/HKU4_15_ in 0.6 M Sodium Chloride, 0.1 M MES-NaOH, pH 6.5 and 20% (w/v) PEG 4000, B6-SARS-CoV/SARS-CoV-2_16_ in 0.2 M Potassium Chloride and 20% (w/v) PEG3350. Crystals were cryoprotected by addition of glycerol to a final concentration of 25% (v/v) and flash cooled in liquid nitrogen. Diffraction data were collected at the beamlines 8.2.1 and 5.0.1 (Advanced Light Source, Berkeley, USA). All data were integrated, indexed and scaled using mosflm (Battye et al., 2011) and Aimless (Evans and Murshudov, 2013) or XDS (Kabsch, 2010). The structures were solved by molecular replacement using Phaser (McCoy et al., 2007) and the S230 Fab (PDB 6NB8) or B6 Fab without ligand as a search model. Model building was performed with Coot (Emsley et al., 2010) and structure refinement with Buster (Blanc et al., 2004) and Phenix (Liebschner et al., 2019). Validation used Molprobity (Chen et al., 2010) and Phenix (Liebschner et al., 2019).

## Data availability

The atomic coordinates and cryoEM maps will be deposited to the Protein Data Bank and Electron Microscopy Data Bank.

